# Genes of the fatty acid oxidation pathway are upregulated in female as compared to male cardiomyocytes

**DOI:** 10.1101/2024.05.29.594850

**Authors:** Maya Talukdar, Lukáš Chmátal, Linyong Mao, Daniel Reichart, Danielle Murashige, Yelena Skaletsky, Daniel M. DeLaughter, Zoltan Arany, Jonathan G. Seidman, Christine Seidman, David C. Page

**Author notes:** These authors contributed equally (Maya Talukdar, Lukáš Chmátal).

## Abstract

Human females and males differ in cardiac physiology and pathology, even after controlling for sex differences in anthropometrics, lifestyle, and environment. For example, females and males differ in cardiac stroke volume and ventricular thickness, and they exhibit different rates and symptoms of cardiovascular disease. Less is understood about molecular differences in female and male hearts, such as sex differences in gene expression. Here we present an integrative framework utilizing bulk and single-nucleus RNA-sequencing data to study sex differences in the cardiac transcriptome. We show that genes of the fatty acid oxidation (FAO) pathway, the primary source of energy in the heart, are expressed more highly in healthy female than in healthy male hearts. We demonstrate that this sex difference is due to cardiomyocyte-specific, female-biased expression of FAO genes and cannot be explained by sex differences in cardiac cellular composition or number of mitochondria, where FAO takes place. Finally, we observe increased cardiac flux and energetic utilization of free fatty acids in female compared to male hearts. Overall, our results demonstrate that male and female human hearts exhibit fundamental differences in metabolism that likely contribute to sex differences in cardiac physiology and pathology.

## Introduction

The heart is a four-chambered pump responsible for the continuous circulation of blood, oxygen, and nutrients throughout the body. Despite this central role in physiology, there are notable differences between human male and female hearts. For example, females have higher heart rates, smaller left ventricles, and reduced stroke volumes compared to males, even when controlling for differences in body size^1,2^. Cardiac pathologies such as myocardial infarctions, cardiomyopathies, and heart failure all display marked sex differences in prevalence and outcome^3–5^. However, the extent to which molecular sex differences, such as sex differences in gene expression, exist between healthy human male and female hearts is largely unknown. Here we analyze several large RNA-sequencing (RNA-seq) datasets of the human heart to study sex differences in the healthy cardiac transcriptome^6–8^. We have previously described significant sex differences in cardiac cellular composition, which could confound efforts to identify and interpret sex differences in gene expression from bulk RNA-sequencing (RNA-seq) data^7^. Thus, we establish a framework combining bulk and single-nucleus RNA-sequencing (snRNA-seq) datasets to identify cell-type-specific sex differences in gene expression *in vivo* (Fig. 1A). Applying this framework to the heart, we find that genes in the fatty acid oxidation (FAO) pathway, the major source of energy in the heart, are more highly expressed in female cardiomyocytes (CMs) compared to male CMs. Using previously reported cardiac metabolomic data, we show that non-failing female hearts exhibit an increased flux and energetic utilization of free fatty acids (FFAs) compared to males. Together, our findings reveal biologically relevant, cell-type-specific sex differences in gene expression, flux, and utilization of the FAO pathway in the human heart.

**Figure 1.**
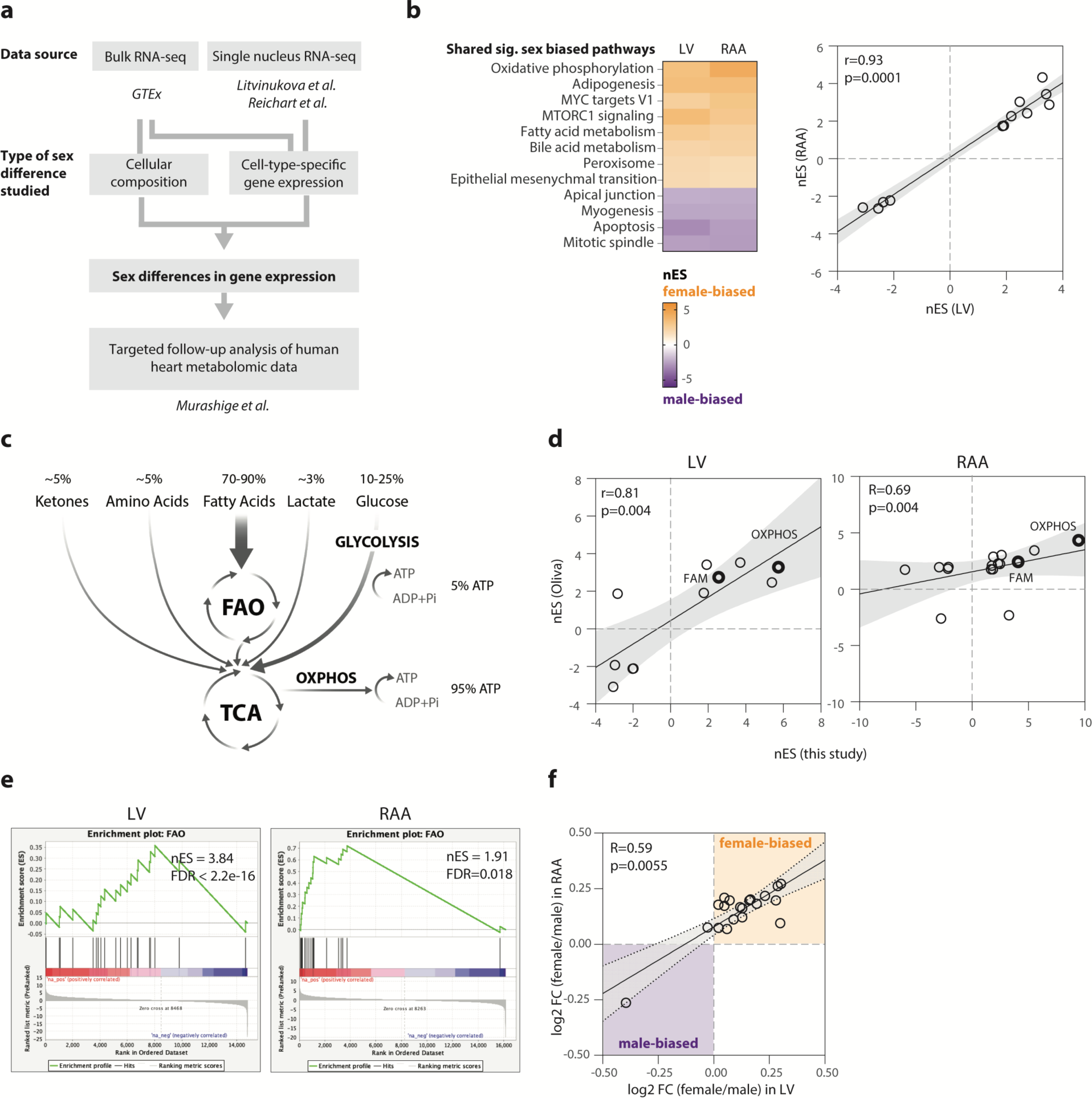
Analysis of GTEx bulk RNA-sequencing data of left ventricle and right atrial appendage reveals many significantly sex-biased pathways, including fatty acid oxidation and oxidative phosphorylation. **(a)** Research design employed here to identify sex differences in healthy human heart. **(b)** Significantly sex-biased (adj. *p* < 0.05) Hallmark Molecular Signature Database pathways identified in both left ventricle (LV) (n = 428 donors) and right atrial appendage (RAA) (n = 429 donors) using gene set enrichment analysis (GSEA) on sex-biased genes identified from Gene Tissue Expression Consortium (GTEx) bulk RNA-sequencing data analyzed in Oliva *et al.*, 2022. These pathways show highly correlated normalized enrichment scores (nES) between LV and RAA. **(c)** Schematic of fuels consumed in cardiac ATP production. **(d)** Spearman correlation of nES of significantly sex-biased pathways mutually identified in this study and Oliva *et al.* in LV and RAA. Oxidative phosphorylation (OXPHOS) and fatty acid metabolism (FAM) are highlighted. **(e)** GSEA running rank plots of fatty acid oxidation (FAO) in LV and RAA. **(f)** Spearman correlation between sex bias of individual FAO genes in LV and RAA. FDR, false discovery rate; FC, fold change.

## Results

To assess sex differences in the cardiac transcriptome, we analyzed bulk RNA-seq data from over 400 donors in version 8 of the Genotype Tissue Expression (GTEx) Project^9^. A previous study of gene expression in 44 GTEx tissues identified more than 1,000 sex-biased genes – most of which were autosomal – in both the left ventricle (LV) and the right atrial appendage (RAA), the two heart regions available in GTEx^6^. However, only a small number of gene sets, all with limited known relevance to cardiac physiology, were found to be significantly enriched among these sex-biased genes. We re-analyzed these previously identified LV and RAA autosomal sex-biased genes using gene set enrichment analysis (GSEA) and the fifty curated Molecular Signature Database Hallmark pathways to further study sex-biased pathways^10,11^. Our analysis identified 12 pathways that were significantly sex-biased in both the LV and RAA, all with highly correlated enrichment scores between these two heart regions and none of which were identified in the original Oliva *et al.*, 2020 analysis (Fig. 1B; Supp. Table 1).

Based on this preliminary finding, we developed a gene expression quantification pipeline with several key innovations to facilitate identification of additional subtle sex biases in expression (Materials and Methods). This included a complete realignment of the GTEx v8 primary RNA-seq data to an updated (hg38) reference genome (Gencode v42). We identified 2,956 and 1,107 sex-biased genes in the LV and RAA, respectively, 417 of which were significantly sex-biased in both regions and most of which were autosomal (Supp. Table 2). Using GSEA and the Hallmark pathways to analyze these autosomal sex-biased genes, we identified a strong concordance of sex-biased pathways using sex-biased genes from our study and Oliva *et al.* (Supp. Table 3). Interestingly, we observed that the most female-biased pathway in the RAA in both studies was oxidative phosphorylation (OXPHOS), which was also female-biased in the LV. OXPHOS produces over 90% of ATP in the healthy heart, primarily by burning fatty acids with additional contributions from glucose metabolism, ketone bodies, and amino acids (Fig. 1C). We observed that the fatty acid metabolism (FAM) pathway was also significantly female-biased in the LV and RAA in both studies (Fig. 1D); we did not observe a significant sex bias for any other metabolic pathway. Given the importance of FAM and OXPHOS for ATP generation in the healthy myocardium, we prioritized these pathways for further investigation^12^.

The Hallmark FAM gene set combines genes involved in fatty acid oxidation (FAO) – and therefore ATP production via OXPHOS – with genes involved in fatty acid synthesis (FAS), which is not a major metabolic pathway in the heart^13^. Accordingly, we performed additional GSEA using curated FAO and FAS gene sets to test if the female-bias of FAM we observed was driven by genes in the FAO pathway^14–16^. We found that the FAO gene set was significantly female-biased in both the RAA and LV while the FAS gene set showed no consistent sex bias (Fig. 1E; Supp. Fig. 1A-B). Further inspection of the individual genes in the FAO pathway revealed female-biased expression across almost all of these genes in both the LV and RAA, with a strong correlation in sex bias fold-change between the two regions (Fig. 1F). We also tested whether the FAO gene set was similarly female-biased in (FAO-dependent) skeletal muscle but observed no significant sex bias, suggesting that FAO gene expression is female-biased specifically in cardiac muscle (Supp. Fig. 1C).

Bulk RNA-seq datasets such as GTEx are invaluable for pursuing well-powered studies of often subtle sex differences in gene expression in human tissues. However, sex differences in cellular composition can confound analyses of bulk RNA-seq data. We previously reported that healthy female hearts have a higher proportion of CMs than healthy male hearts^7^. As CMs are the primary contractile cells of the heart and therefore highly reliant on ATP derived from FAO via OXPHOS, we reasoned that the female-bias of FAO and OXPHOS we observed in bulk RNA-seq data could reflect sex differences in CM abundance, or cell-type-specific sex differences in FAO and OXPHOS expression, or both. To distinguish among these possibilities, we analyzed cardiac single-nucleus RNA-seq (snRNA-seq) data to specifically identify cell-type-specific sex differences in gene expression. We integrated two large human snRNA-seq datasets we previously published to create an atlas of >450,000 high-quality heart nuclei from 12 male and eight female middle-aged donors with no known cardiac pathology (Fig. 2A; Supp. Figs. 2-3). These nuclei were isolated from tissues from all four chambers of the heart – the left atrium (LA), left ventricle (LV), right atrium (RA), and right ventricle (RV) – as well as the septum (SP) and apex (AX). Using our atlas, we identified all major cardiac cell types and confirmed that females harbored a higher proportion of LV CMs than males (adj. *p =* 0.024, two-tailed t-test), with no other regional cell type showing a significant sex difference in proportion (Fig. 2B-C; Supp. Figs. 4-5).

**Figure 2.**
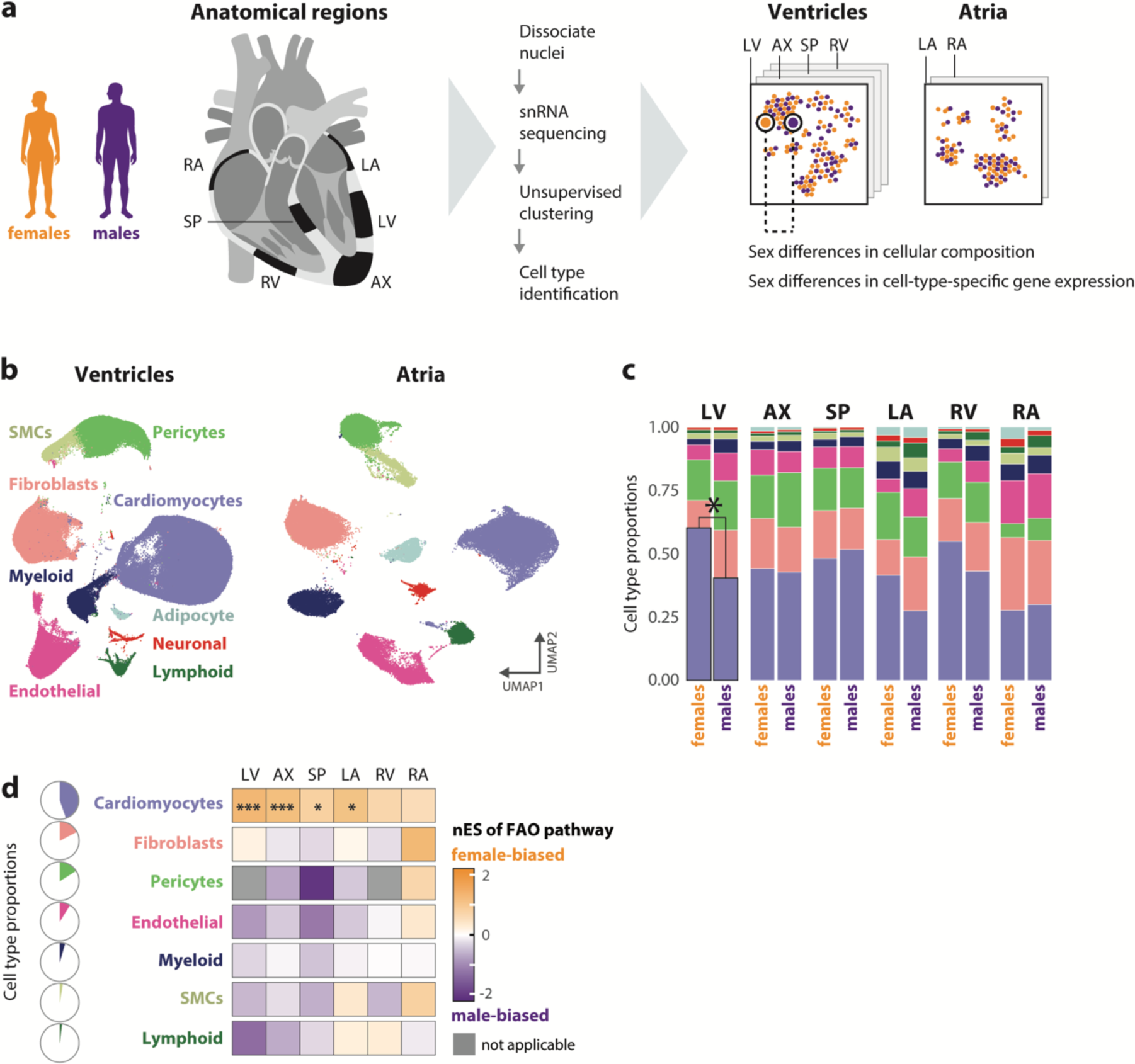
Female bias in FAO gene expression is specific to cardiomyocytes in all heart chambers and is not driven by sex differences in cardiac cell type proportions. **(a)** Schematic of processing and analytical pipeline for female and male healthy heart samples (n = 8 and n = 12, respectively) collected from distinct anatomical regions and used in this study. LV, left ventricle (n = 19); AX, apex (n = 13); SP, septum (n = 17); LA, left atrium (n = 12); RV, right ventricle (n = 16); RA, right atrium (n = 12). **(b)** UMAP embeddings of ventricular and atrial single-nucleus RNA-seq data. (**c**) Proportions of cardiac cell types in female and male anatomical regions. Only LV cardiomyocytes show a significant sex difference in proportion (two-tailed t-test, adj. *p* = 0.026). (**d**) Sex bias of FAO pathway using gene set enrichment analysis across all cardiac regions and cell types represented by at least 5,000 nuclei. Significant female-biased (orange) and male-biased (purple) pathways: * adj. *p* < 0.05, ** adj. *p* < 0.01, *** adj. *p* < 0.001. SMCs, smooth muscle cells; nES, normalized enrichment score.

The breadth of our integrated atlas allowed us to identify genes whose expression is sex-biased in a specific cell type in one or more regions of the heart (Materials and Methods). Overall, we identified 8,167 protein-coding genes with significant sex-biased expression in at least one regional cell-type of the heart; 7,918 (97%) of these genes are autosomal (Supp. Table 4). Using downsampling analysis, we confirmed that we are well-powered to identify sex-biased genes for nearly all regional cell-types (Materials and Methods; Supp. Figs. 6-7). Thus, we next used our integrated snRNA-seq atlas to dissect the female-biased FAO and OXPHOS gene expression we observed in bulk RNA-seq data. Using GSEA to analyze autosomal sex-biased genes, we found that the FAO pathway was significantly female-biased in LV, SP, AX, and LA CMs (Fig. 2D; Supp. Table 5). We also observed that FAO was female-biased in RA and RV CMs, although this did not reach statistical significance. As we were similarly powered to identity sex-biased genes in the left and right heart chambers, this stronger female bias in LV, SP, AX, and LA CMs may reflect an increased energetic demand of the left heart, which is responsible for the high-pressure systemic circulation, compared to the right heart, which is responsible for the low-pressure pulmonary circulation. Interestingly, no other cell type showed a significant sex bias in FAO or a consistent trend across the six heart regions we assessed. This includes cell types like myeloid cells and fibroblasts that also utilize FAO for energy production (Supp. Fig. 8). Inspection of each gene within the FAO pathway revealed strikingly concordant sex bias effect sizes across all regional CMs, with 18 out of 20 FAO genes expressed approximately 10 – 30% higher in female CMs compared to male CMs (Fig. 3A-B). This set of consistently female-biased genes encompassed all major enzymes of the cardiac FAO pathway, including *CPT1B*, which catalyzes the rate-limiting step of FAO in the heart and serves as a major nexus of metabolic regulation in CMs^17^. Moreover, we reasoned that a biologically-relevant female bias in expression of FAO genes would be accompanied by female-biased expression of plasma membrane-bound fatty acid transporters (FATs), which transport fatty acids from the blood to the cytoplasm. Consistently, we found that expression levels of most FATs were higher in females, although effect sizes were more modest than for the core FAO pathway enzymes (Fig. 3C). We observed a particularly striking female bias of FAT *CD36*, which was female-biased in five of the six cardiac regions assessed^18^. We also investigated expression of the OXPHOS pathway, where unlike FAO we observed no directionally consistent sex bias across heart regions (Supp. Fig. 9).

**Figure 3.**
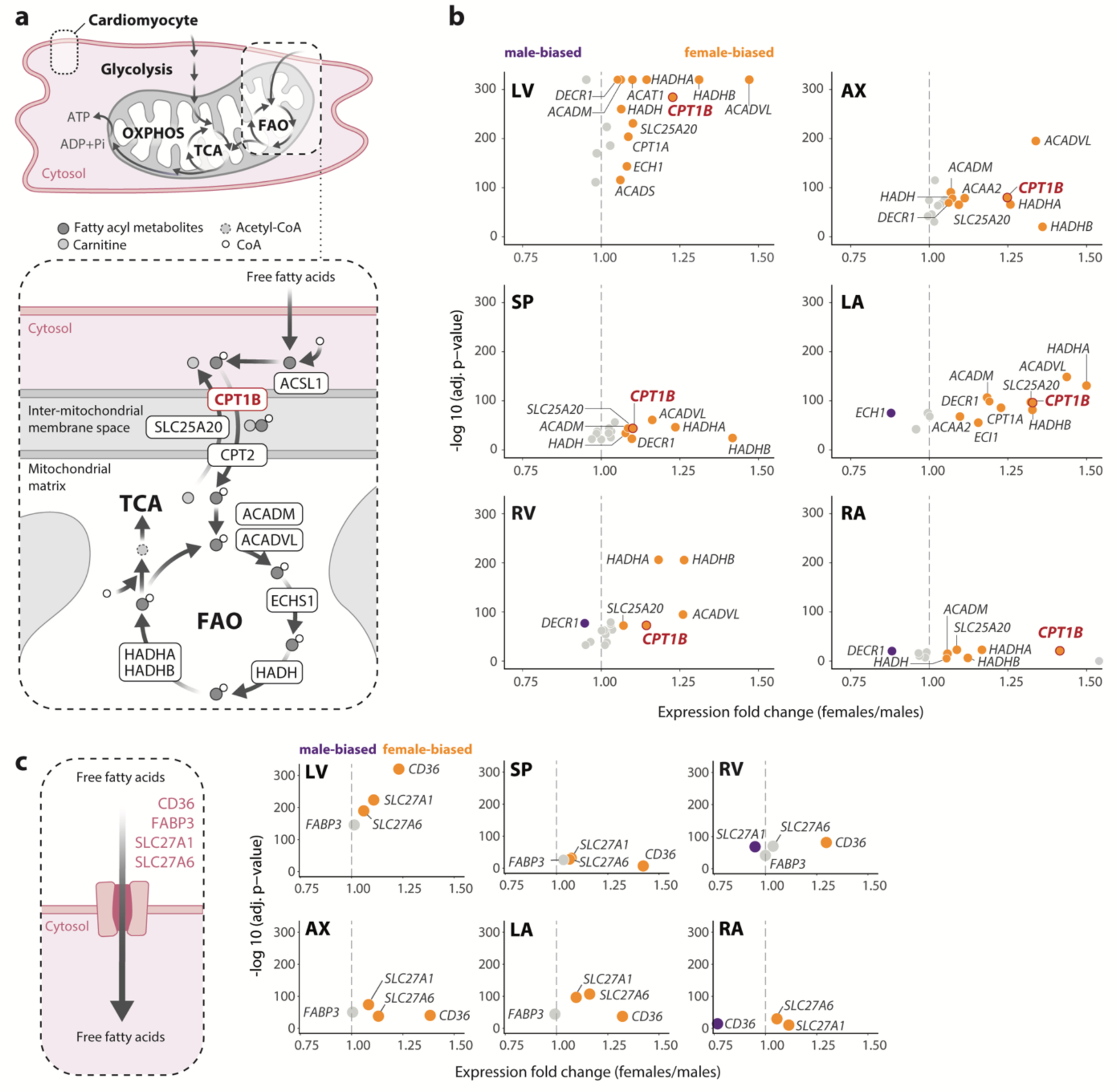
FAO pathway genes are more highly expressed in female cardiomyocytes in all heart regions. **(a)** Schematic of cardiomyocyte energy metabolism, highlighting the FAO pathway. *CPT1B*, the rate-limiting enzyme of FAO, in red. **(b)** Volcano plots of FAO genes significantly differentially expressed between male (purple; FC < 0.95, adj. *p* < 0.05) and female cardiomyocytes (orange, FC > 1.05, adj. *p* < 0.05) in each heart region, with non-significant genes in gray. **(c)** Volcano plots of fatty acid transporter genes significantly differentially expressed between male (purple; FC < 0.95, adj. *p* < 0.05) and female cardiomyocytes (orange, FC > 1.05, adj. *p* < 0.05) in each heart region, with non-significant genes in gray. OXPHOS, oxidative phosphorylation; TCA, tricarboxylic acid cycle; CoA, coenzyme-A.

Given that FAO and glycolysis are two major sources of fuel for the healthy heart, we reasoned that a female bias in expression of FAO genes might be compensated for by male-biased expression of glycolysis genes. Indeed, a previous study that performed single-nucleus ATAC-seq on healthy male and female hearts reported that male-biased chromatin accessibility peaks are enriched for motifs of transcription factors that upregulate the glycolytic pathway^20^. However, we did not observe such a trend in our data (Supp. Fig. 10); on balance, then, there is insufficient evidence to infer increased use of glycolysis in the male heart, at least as mediated by transcriptional mechanisms. We also investigated whether expression of genes of the tricarboxylic acid cycle is sex-biased in CMs but again did not see a consistent trend across all regions (Supp. Fig. 11).

Our snRNA-seq analysis suggests that expression of FAO genes is female-biased specifically in CMs. To further pursue this finding, we investigated the alternative hypothesis that proportions of CM subpopulations differ between female and male hearts. We previously reported and validated several CM subpopulations within both the atria and ventricles, including subpopulations defined by metabolic characteristics^7^. Similar to the potential influence of sex differences in cell-type composition in bulk RNA-seq data, sex differences in abundance of these CM subpopulations could confound our observed female-biased expression of FAO genes in CMs. To assess this possibility, we first asked whether there were significant differences in proportions of CM subpopulations between females and males; we found no such differences (Fig. 4A). We next performed sex-biased differential expression analysis within each CM subpopulation and confirmed significant female-biased expression of FAO in nearly all subpopulations (Supp. Tables 6-7). We conclude that the higher expression of FAO genes we observe in female CMs is not due to sex differences in proportions of CM subpopulations (Fig. 4B).

**Figure 4.**
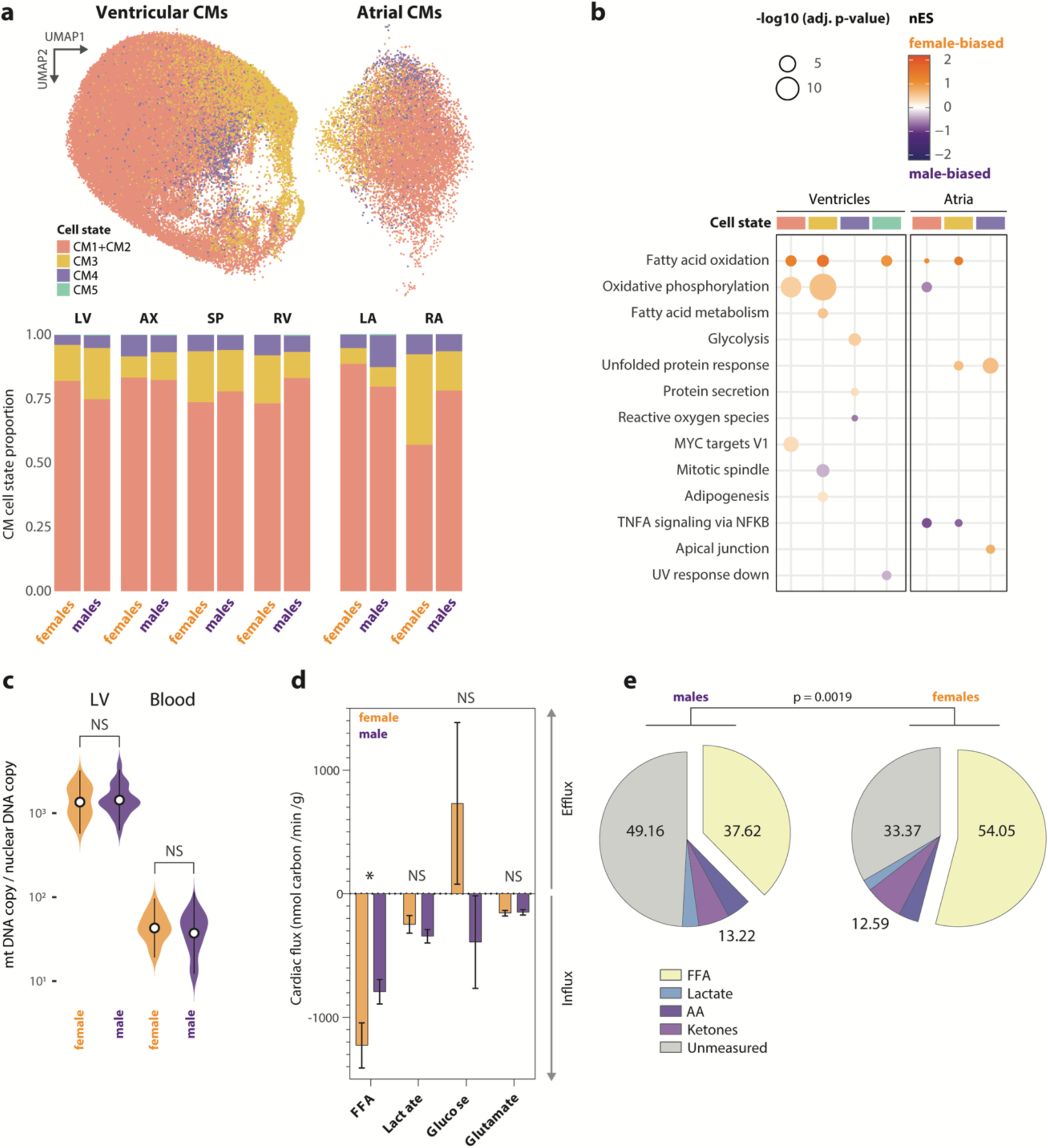
Female-biased expression of FAO in cardiomyocytes cannot be explained by sex differences in mitochondrial content or cardiomyocyte heterogeneity and is mirrored by sex differences in cardiac free fatty acid flux. **(a)** UMAP embeddings of snRNA-seq data with cardiomyocyte cell states and cell state proportions in each sex and cardiac anatomical region. **(b)** Gene set enrichment analysis shows significantly sex-biased pathways in each CM regional cell state (adj. *p* <0.05). **(c**) Relative mitochondrial copy number in non-diseased female (orange; n = 12) and male (purple; n = 20) LV samples from GTEx and corresponding whole blood samples from the same individuals**. (d)** Myocardial fluxes of free fatty acids (FFA), lactate, glucose, and glutamate (median values with 95% confidence intervals) in non-failing female (orange; n = 34) and male hearts (purple; n = 53) calculated with data from Murashige *et al*., 2020 and Ngo *et al.,* 2022. Only FFA shows a significant sex difference in myocardial flux (Welch’s t-test, *p* = 0.037). **(e)** Proportional contributions of classes of metabolites to maximal oxygen consumption in male and female myocardium calculated with data from Murashige *et al.* (chi-squared proportion test, two-tailed *p* = 0.0019). OXPHOS, oxidative phosphorylation; TCA, tricarboxylic acid cycle; NS, not significant. LV, left ventricle; AX, apex; SP, septum; RV, right ventricle; LA, left atrium; RA, right atrium; CMs, cardiomyocytes; nES, normalized enrichment score.

While FAO genes are encoded in the nucleus, the enzymatic reactions of the FAO pathway take place in the mitochondria. Thus, sex-biased FAO gene expression could be driven by sex differences in mitochondrial copy number (mtCN), as a higher abundance of mitochondria might require overall increased nuclear expression of FAO genes even if expression of FAO genes within each mitochondrion is not sex-biased. Accordingly, we tested the hypothesis that female-biased expression of FAO genes within CMs is due to increased mtCN in the female heart. We obtained 20 male and 12 female non-diseased LV and matched whole blood samples from GTEx, and we isolated genomic DNA and performed quantitative PCR to calculate relative mtCN from genomic DNA, as previously described (Materials and Methods)^21^. Reassuringly, we observed a significantly higher mtCN in LV versus whole blood across donors but no significant sex differences in either blood or LV mtCN (Fig. 4C), which is consistent with prior reports^22^. We conclude that the higher expression of FAO genes we observe in female CMs is not due to sex differential mtCN.

We next sought to understand the biological relevance of this female-biased CM FAO gene expression to overall cardiac metabolism. We reasoned that higher expression of FAO core enzymes and FATs predicts higher fatty acid utilization by the female heart versus the male heart. To test this prediction, we re-analyzed cardiac metabolomic data obtained from the radial artery and coronary sinus of 34 female and 53 male individuals who presented for elective catheter ablation of atrial fibrillation and had no history of heart failure or reduced ejection fraction^12^. Cardiac flux is calculated as the product of 1) measured cardiac uptake per metabolite, considering the metabolite’s concentration and carbon composition, and 2) myocardial blood flow, which varies with sex and age. Thus, we calculated cardiac flux for each donor, incorporating previously published age-and sex-specific myocardial blood flow values measured from 1,463 healthy individuals (Materials and Methods)^23^. We found that free fatty acids (FFAs) showed similar arterial concentrations but female-biased cardiac uptake, with significantly higher cardiac flux (fold-change = 1.55; *p* = 0.038, Welch’s t-test) in females than in males (Materials and Methods; Fig. 4D; Supp. Fig. 12). We did not observe a significant sex difference in the cardiac flux of any other metabolite, including lactate, glucose, and amino acids.

Finally, we calculated the proportional contribution of various metabolites to total ATP production in the heart. Oxygen consumption was measured for a subset of patients (ten males and seven females) to determine total cardiac ATP requirement, and the energetic contribution of measured metabolites was calculated assuming full oxidation. The difference between total ATP consumption and the ATP produced by full oxidation of measured metabolites can be attributed to metabolites that were not assayed, including glucose and lipoprotein-bound fatty acids. There was a significant sex difference in proportional use of fuels for cardiac ATP production (chi-squared proportion test, *p* = 0.0019; Fig. 4E), with FFAs providing approximately 54% of ATP in the female heart and 38% of ATP in the male heart. Thus, while future work will be required to assess sex differences in total fatty acid consumption (including both free and lipoprotein-bound FAs), our data is consistent with a higher utilization of FFAs in the female as compared to male heart. Taken together with our transcriptomic analyses, we establish that the healthy human female heart exhibits higher expression and flux through the FAO pathway as well as increased energetic reliance on FFA oxidation compared to the male heart.

## Discussion

We find that healthy human female cardiomyocytes express FAO genes more highly than male cardiomyocytes, and that this result cannot be explained by sex differences in cardiomyocyte subpopulations or mitochondrial copy number. Importantly, this molecular sex difference can be identified from bulk RNA-sequencing data but can only be directly attributed to cardiomyocyte expression through tandem analysis of snRNA-seq data, presenting a new paradigm for studying subtle molecular sex differences using complementary types of transcriptomic data. We further show that the nonfailing female heart experiences a significantly higher flux of FFAs – but not lactate, glucose, or amino acids – as well as higher proportional energetic consumption of FFAs as compared to the male heart.

Further studies will be required to identify the mechanisms driving the female-biased expression and flux of the FAO pathway in the heart, including identifying the relative contributions of sex hormones and sex chromosomes. For example, many genes in the FAO pathway are transcriptionally activated by *PPARA* and *PPARGC1A*, which also interact with estrogen-related receptors and were generally female-biased in our RNA-seq analysis (Supp. Fig. 13)^24^. Consistently, prior work in rats has established that estrogen replacement can stimulate cardiac FAO even in ovariectomized female rats, highlighting the potential role of estrogen in shaping female-biased expression of FAO genes^25^. Studies in mice have found that sex chromosome complement influences general adiposity and lipid metabolism, independent of gonadal sex^26–28^. As the sex differences in FAO we observe in humans are present when comparing likely post-menopausal donors (age > 55) to age-matched males, our analyses additionally support a sex-chromosome-dependent mechanism that explains how female-biased expression of FAO genes can be maintained even as estrogen levels decrease with age.

While this report focuses on the non-diseased human heart, we anticipate that the observed sex differences in metabolism will inform our understanding of sex differences in cardiac disease. Most cardiac diseases show sex differences in prevalence and outcomes, and nearly all cardiac diseases involve dysregulated heart metabolism, including altered lipid metabolism^29,30^. Autosomal disorders of FAO often present with cardiac phenotypes such as dilated cardiomyopathy that are associated with worse outcomes and higher mortality in affected males compared to affected females^31,32^. Understanding how baseline sex differences in FAO interact with disease-associated metabolic shifts will be crucial to understanding sex-biased heart disease pathogenesis. Overall, we report female-biased expression and flux of the FAO pathway – a novel, cell-type-specific, and physiologically-relevant molecular sex difference in the human heart.

## Materials and Methods

### Processing and alignment of GTEx data

A bam file for each GTEx (v8) RNA-sequencing sample was downloaded from the AnVIL repository, and Picard (v2.23.3) was used to convert the bam file to paired-end (PE) FASTQ files. STAR (v2.7.1) was then used to map PE reads to the GRCh38 human reference genome. After alignment, htseq-count (v1.99.2) was used with the parameter setting “--mode=union --nonunique=none” to estimate numbers of read pairs (i.e., fragments) that derived from each gene. Thus, only uniquely mapped fragments were used. In addition, fragments that align to or overlap with more than one gene were excluded. The Gencode v42 gene annotation was used when counting read pairs per gene.

### Sex-biased gene expression analysis of GTEx data

For each tissue, fragment counts were normalized using edgeR (v3.34.1) and lowly-expressed genes (median counts per million per tissue < 0.5) were filtered out. Sex-biased genes were identified with edgeR per tissue, with age and BMI as covariates.

### Gene set enrichment analysis of GTEx data from Oliva *et al*., and this study

Gene set enrichment analysis (GSEA) was performed using the GSEA-MSigDb software released by the Broad Institute (v4.1.0; command-line version). Autosomal genes were ordered from most female-biased to most male-biased genes for both Oliva *et al.* and the current study’s differential expression results. GSEA-MSigDb was run with an unweighted scoring scheme (“-scoring_scheme classic”) against the 50 Hallmark Molecular Signature Database categories, as well as fatty acid oxidation (FAO) and fatty acid synthesis (FAS) gene sets. The FAO gene set was extracted from Houten et al., 2016, which includes FAO genes that are primarily expressed in cardiac tissue (ex: CPT1B). The FAS gene set was extracted from the gene ontology biological processes (GO BP; GO:0006633) pathway.

### Alignment and expression quantification from single-nucleus RNA-sequencing FASTQ files

Alignment of single-nucleus RNA-sequencing (snRNA-seq) FASTQ files was performed using CellRanger (v7.1.0) and CellRanger’s pre-built human (hg38) reference genome. --include-introns and all other default parameters were used.

### Quality control and pre-processing of snRNA-seq data

CellBender (v0.30.0) was used with default parameters to remove background ambient RNA contamination, and scrublet (v0.2.3) was used to detect and remove putative doublets on each sample individually. Atrial and ventricular samples were then merged into two Seurat (v4.4.0) objects, respectively, and nuclei that met the following quality control metrics were retained: 1) percent mitochondrial gene expression < 5%; 2) percent ribosomal gene expression < 5%; 3) 6000 > number of expressed genes per nucleus (nFeature) > 300; and 4) 25,000 > per-nucleus counts (nCount) > 300.

### Cell type/state annotation and sex-biased differential abundance analysis of snRNA-seq data

Cell types were annotated within both the atrial and ventricular Seurat objects using canonical marker genes for adipocytes (*GPAM*, *FASN*, *LEP*), atrial cardiomyocytes (*NPPA*, *MYL7*, *MYL4*), endothelial cells (*VWF*, *PECAM1*, *CHD5*), fibroblasts (*DCN*, *PDGFRA*), lymphoid cells (*CD8A*, *IL7R*, *CD40LG*), myeloid cells (*CD14*, *C1QA*, *CD68*), neurons (*NRXN1*, *NRXN3*, *PLP1*), pericytes (*RGS5*, *ABCC9*, *KCNJ8*), smooth muscle cells (*MYH11*, *TAGLN*, *ACTA2*), and ventricular cardiomyocytes (*MYH7*, *MYL2*, *FHL2*). Cell states were extracted from the original source publications Litvinukova et al., 2020 and Reichart et al., 2022. aCM1/aCM2 and vCM1/vCM2 were merged due to their similar transcriptional profiles. Regional cell types and cardiomyocyte cell states were each tested for differential abundance between male and female donors using a t-test with a Benjamini-Hochberg correction for multiple hypothesis testing.

### Sex-biased gene expression analysis of snRNA-seq data

Sex-biased genes were identified for each regional cell type represented by at least 5,000 nuclei as well as each cardiomyocyte cell state using the Seurat (v4.4.0) function FindMarkers and MAST (v1.28.0), which employs a linear mixed model approach that allows for differential expression analysis while controlling for correlations between individual cells from the same donor (“test.use = ‘MAST’”; “latent.vars = c(‘Donor_Name’)”). Only genes expressed in >10% of cells (“min.pct = 0.10”) were assessed and no average log-2 fold-change was imposed (“avg_log2FC = 0”). All other default parameters were employed.

### Gene set enrichment analysis of snRNA-seq data

GSEA was performed using the limma (v3.58.1) function geneSetTest with all default parameters and the gene sets from the analysis of GTEx bulk RNA-sequencing data; namely, the 50 Molecular Signature Database Hallmark pathways, a FAO gene set (Houten et al., 2016), and a FAS gene set (GO:0006633). Multiple hypothesis testing correction was performed using a Benjamini-Hochberg correction.

### Downsampling analysis of sex-biased genes and pathways in snRNA-seq data

snRNA-seq data was randomly downsampled to 10%, 20%, 30%, 40%, 50%, 60%, 70%, 80%, and 90% (5 random samples per downsampling percentage) for each regional cell type. Sex-biased gene expression analysis and GSEA, as described above, were performed per down sample to determine if sex-biased genes and pathways were saturated at this current snRNA-seq atlas. Most regional cell types showed saturation of sex-biased pathways and genes.

### Analysis of mitochondrial copy number in GTEx left ventricle and whole blood samples

Human heart left ventricle samples (20 males, 12 females) with no medical record of heart disease, as assessed from GTEx metadata and histological evaluation by a cardiac pathologist, were obtained from GTEx. Genomic DNA, including mitochondrial DNA, from heart tissues was isolated using Genomic DNA isolation kit (QIA Amp DNA Mini Kit 50 cat #51304) using ∼ 5 mg of the heart tissue. Relative mitochondrial copy number (mtCN) was analyzed by quantitative PCR (qPCR) using primers for the mitochondria-encoded gene MT-ND2 (forward primer: 5′–TGTTGGTTATACCCTTCCCGTACTA–3′; reverse primer: 5′–CCTGCAAAGATGGTAGAGTAGATGA–3′) and nuclear-encoded gene BECN1 (forward primer: 5′–CCCTCATCACAGGGCTCTCTCCA–3′; reverse primer: 5′– GGGACTGTAGGCTGGGAACTATGC–3′), as described previously (Wanet et al., 2014). All primers were tested for efficiency (E MT-ND2 = 0.923; E BECN1 = 0.921) and linearity (R2 MT-ND2 = 0.99; R2 BECN1 = 1). Heart genomic DNA samples were subsequently analyzed in triplicates using Power SYBR Green PCR Master Mix (Thermo Fisher Scientific) according to the manufacturer’s protocol on a 7500 Fast Real-Time PCR System (Applied Biosystems). mtCN in male and female heart samples was calculated as 2 x 2 (Ct^BECN1^ – Ct^MT-ND2^) and analyzed using a two-tailed t-test, as previously described (Wanet et al., 2014). Blood genomic DNA samples were obtained from the GTEx biobank and corresponded to the same individuals as the left ventricle samples. mtCN in genomic DNA from blood was calculated, following the same protocol used for the heart left ventricle samples.

### Calculation of cardiac metabolic flux

Average carbon flux in nmol carbon ⁎ min^-1^ ⁎ g^-1^ per metabolite was calculated using the following two metrics: 1) measured carbon cardiac uptake (nmol * number of carbon atoms in metabolite), and 2) estimated resting myocardial blood flow (min^-1^ ⁎ g^-1^). Measured carbon cardiac uptake ((C_coronary sinus_ – C_coronary artery)_ * number of carbon atoms) was determined per metabolite using data from 34 females and 53 males undergoing voluntary cardiac ablation for atrial fibrillation (Murashige *et al*., 2020). Resting myocardial blood flow values of healthy individuals, stratified by age and sex, were obtained from *Ngo et al.*, 2022. Average carbon flux was calculated for each individual in Murashige *et al*., and Welch’s t-test was used to compare average carbon fluxes per metabolite between males and females. When calculating average carbon flux, the functionally related free fatty acid metabolites C18:1, C18:2, and C16:0 were combined.

### Calculation of metabolite contribution to absolute cardiac ATP consumption

Metabolite contribution to absolute ATP consumption in 53 male and 34 female individuals was calculated using the data and methodology established in Murashige *et al.*, 2020. In brief, the total cardiac ATP requirement ("total ATP consumption") for male and female donors was calculated by measuring blood oxygen concentration in 10 male and 7 female individuals. The energetic contribution of measured metabolites ("measured ATP consumption") in Murashige *et al*.’s assay was calculated across all donors and assumes full oxidation per metabolite (-1 * average metabolite consumption (uM) * 0.6 * ATP yield for each molecule of a given metabolite). The contribution of unmeasured metabolites such as lipoprotein-derived fatty acids ("unmeasured ATP consumption") to total ATP consumption was calculated as total ATP consumption-measured ATP production. A chi-square test was used to test differences in the proportional contribution of metabolites to total ATP consumption between male and female hearts.

---

**Supplementary Figure 1.**
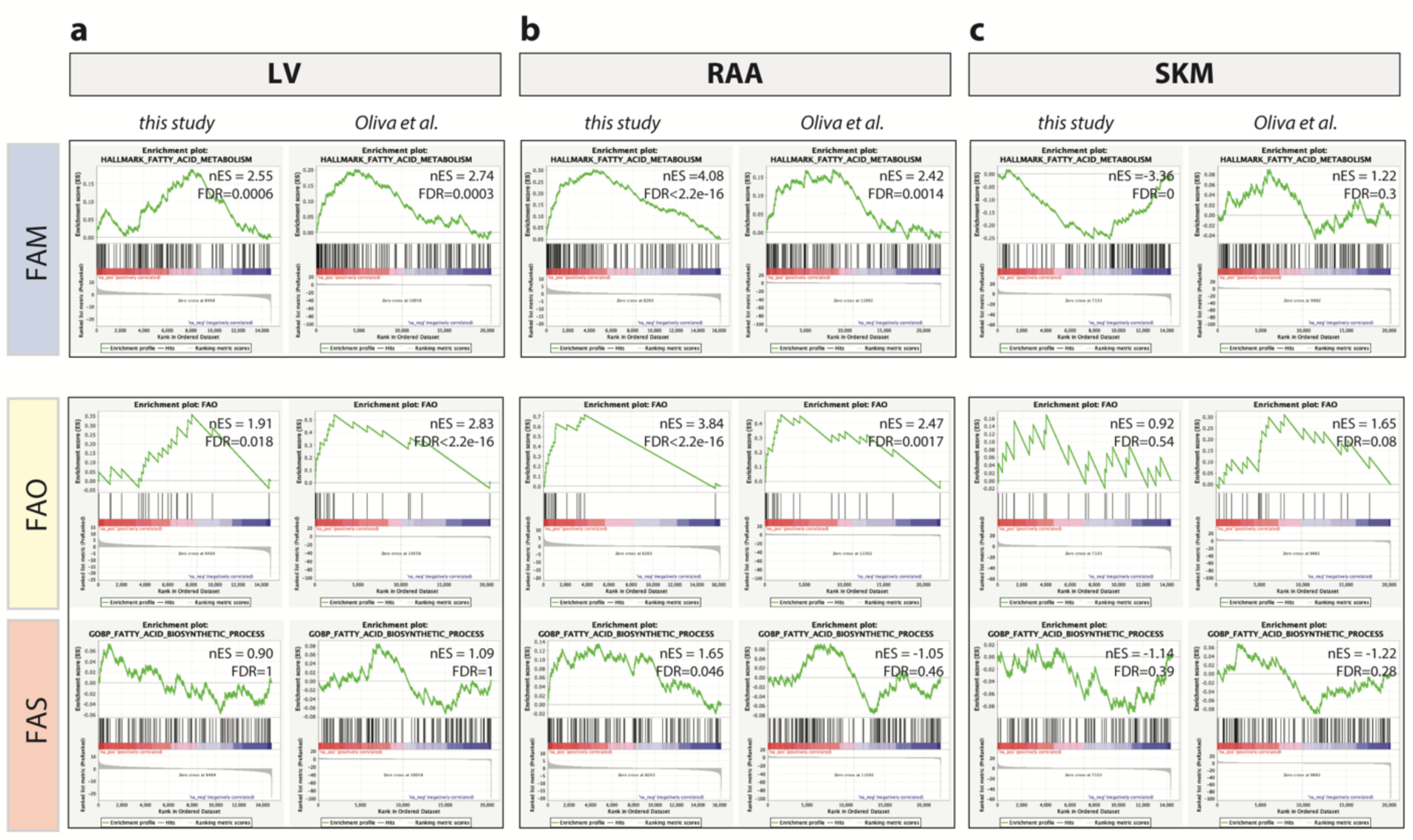
Genes involved in fatty acid oxidation display female-biased expression in GTEx heart tissues but not in skeletal muscle. Running rank plots of gene set enrichment analysis performed for fatty acid metabolism (FAM; from Hallmark Molecular Signatures database), fatty acid oxidation (FAO; from Houten *et al.*, 2016), and fatty acid synthesis (FAS; from GO:0006633). This analysis was performed on sex-biased genes identified in GTEx from this study and Oliva *et al.* across three tissues: **(a)** left ventricle (LV), **(b)** right atrial appendage (RAA), and **(c)** skeletal muscle (SKM).

**Supplementary Figure 2.**
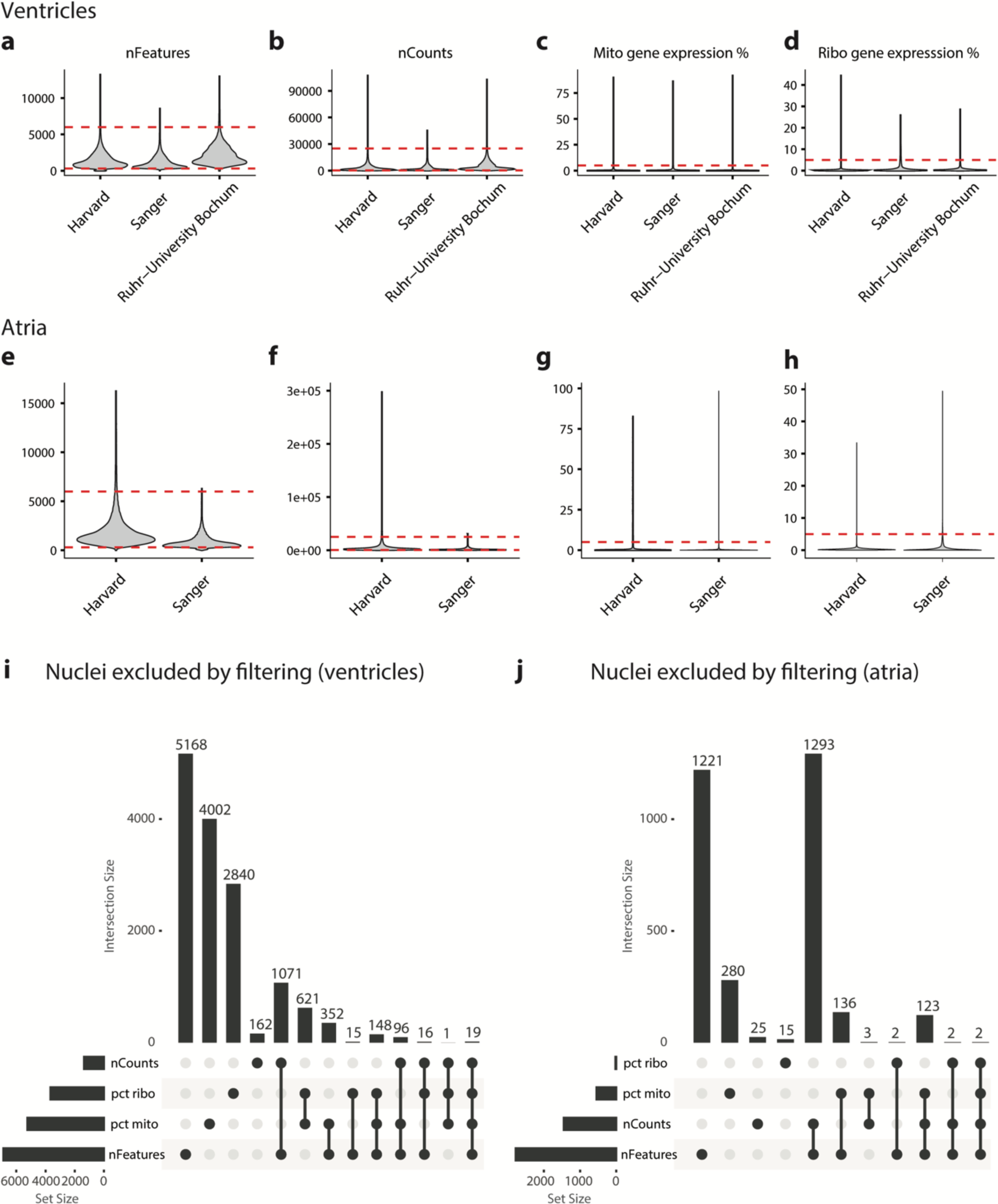
Quality control metrics for single-nucleus RNA-sequencing atlas. **(a-d)** Ventricular and **(e-h)** atrial per-nucleus quality control metrics and filtering thresholds implemented in this study (red dashed lines), split by institution where data was generated. **(i, j)** UpSet plots for **(i)** ventricles and **(j)** atria displaying numbers of nuclei removed based on various combinations of filtering metrics. nFeatures, numbers of expressed genes per nucleus; nCounts, number of unique molecular identifiers per nucleus; pct mito, percentage of mitochondrial gene expression; pct ribo, percentage of ribosomal gene expression.

**Supplementary Figure 3.**
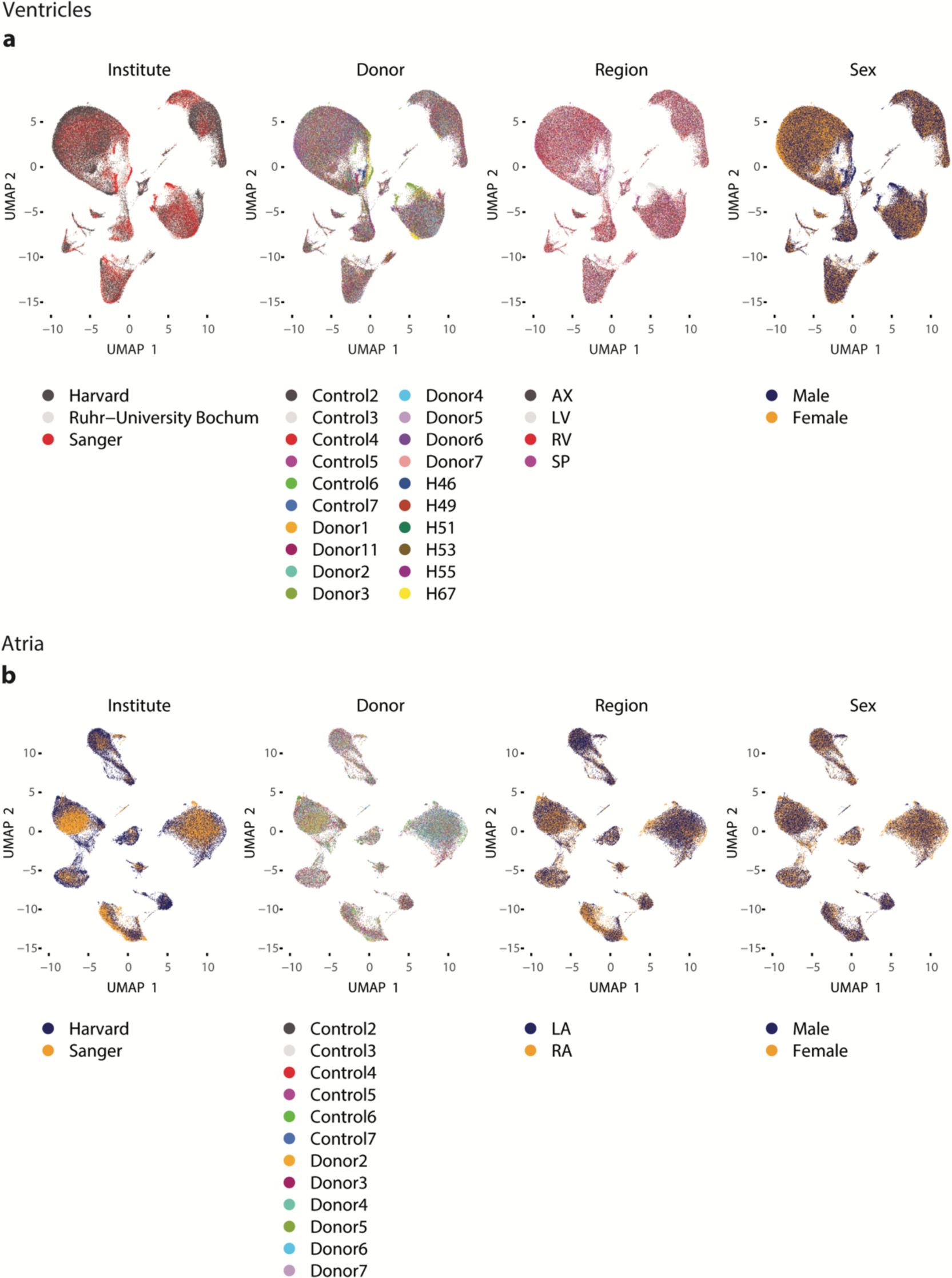
Integration of single-nucleus RNA-sequencing atlas. UMAPs of **(a)** ventricles and **(b)** atria showing successful integration over the covariates of 1) institution where data was generated, 2) donor, 3) cardiac region, and 4) sex. LV, left ventricle; AX, apex; SP, septum; LA, left atrium; RA, right atrium; RV, right ventricle.

**Supplementary Figure 4.**
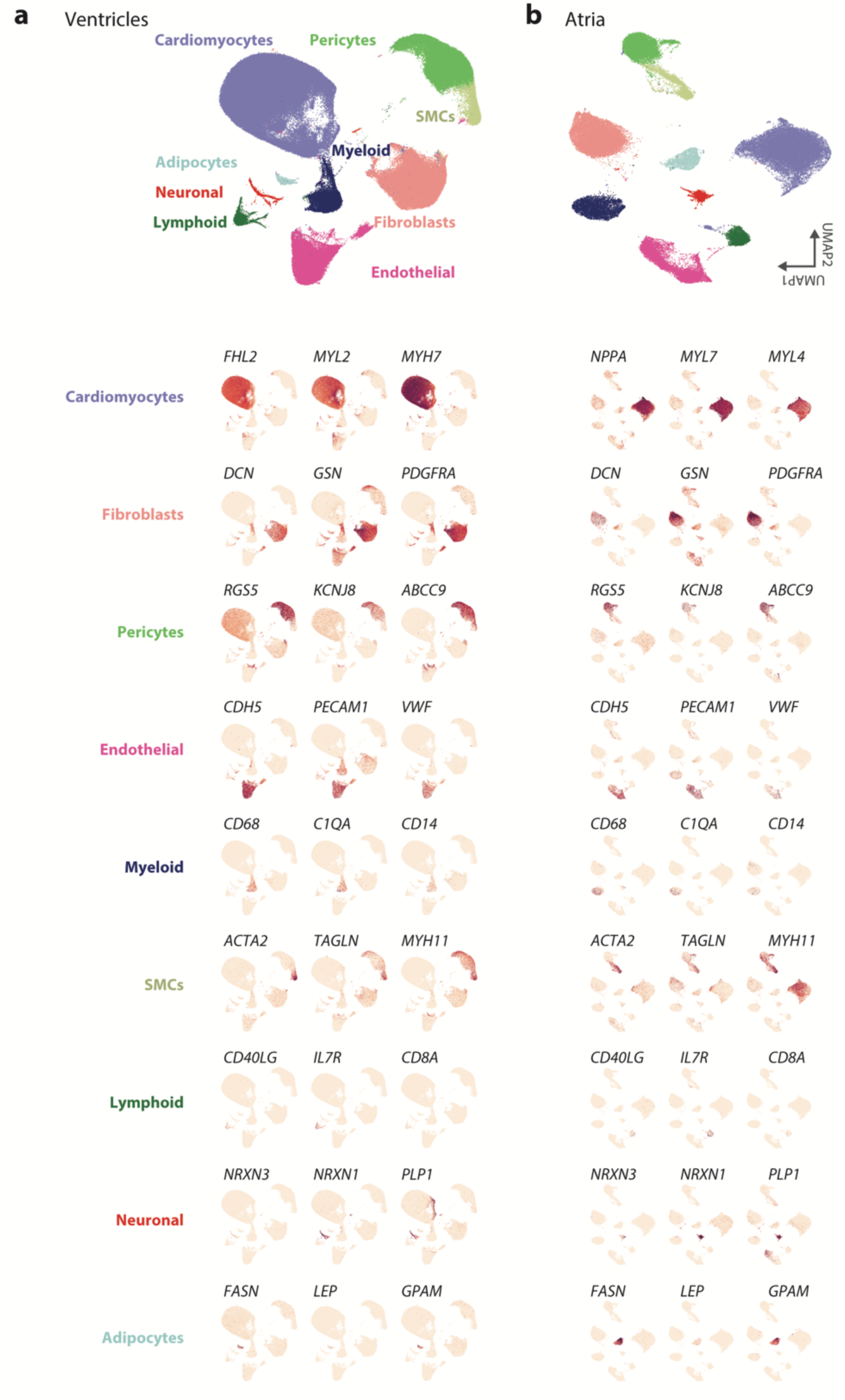
Marker gene visualization for cell type annotation in single-nucleus RNA-sequencing atlas. UMAP embeddings of all cardiac cell types in **(a)** ventricles and **(b)** atria with visualization of canonical cell type markers.

**Supplementary Figure 5.**
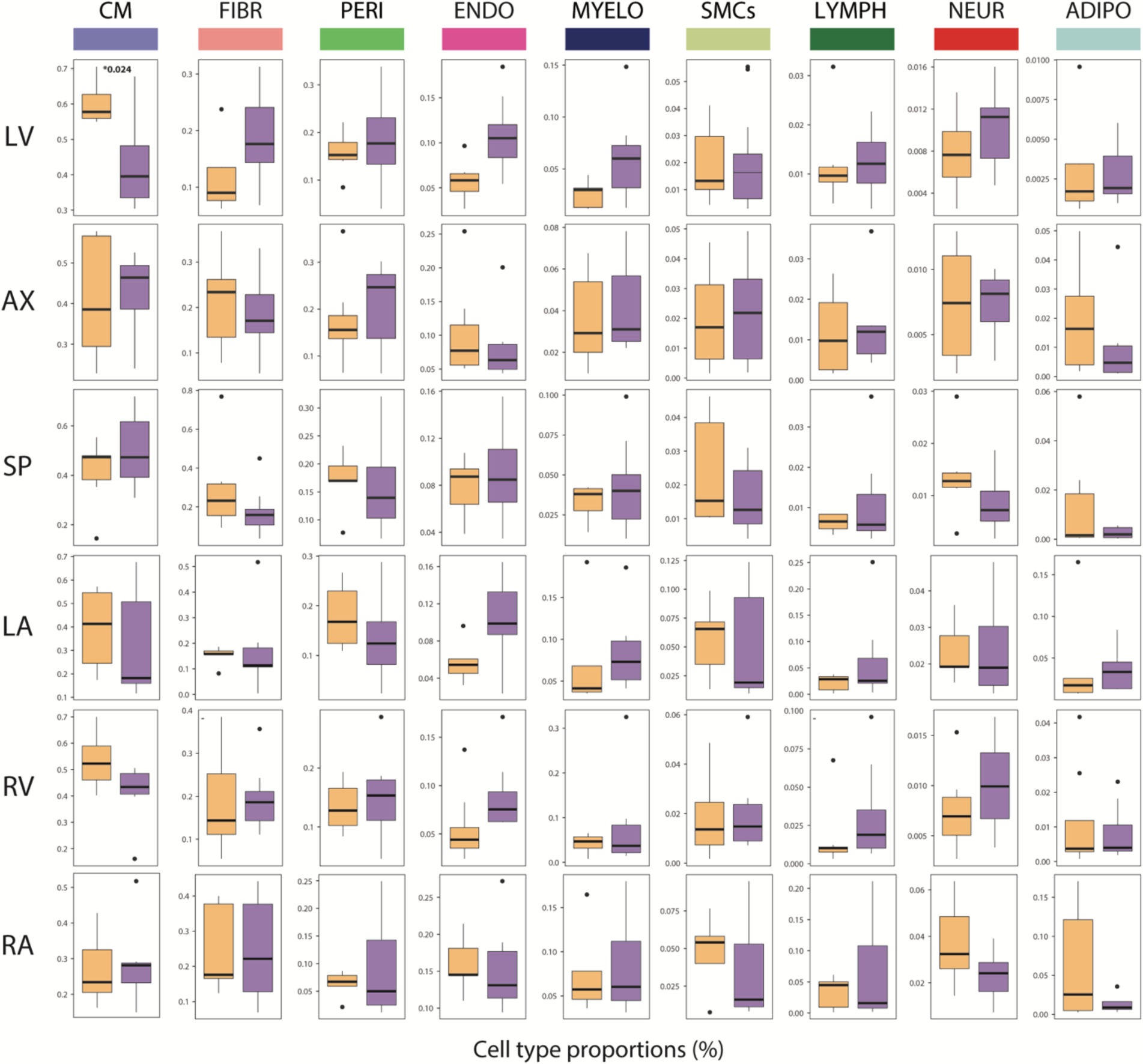
Sex differences in proportions of cell types across heart regions. Two-tailed t-test, adj. *p* < 0.05 (*). CM, cardiomyocytes; Fibr, fibroblasts; Peri, pericytes; Endo, endothelial cells; Myelo, myeloid cells; SMCs, smooth muscle cells; Lymph, lymphoid cells; Neur, neuronal cells; Adipo, adipocytes.

**Supplementary Figure 6.**
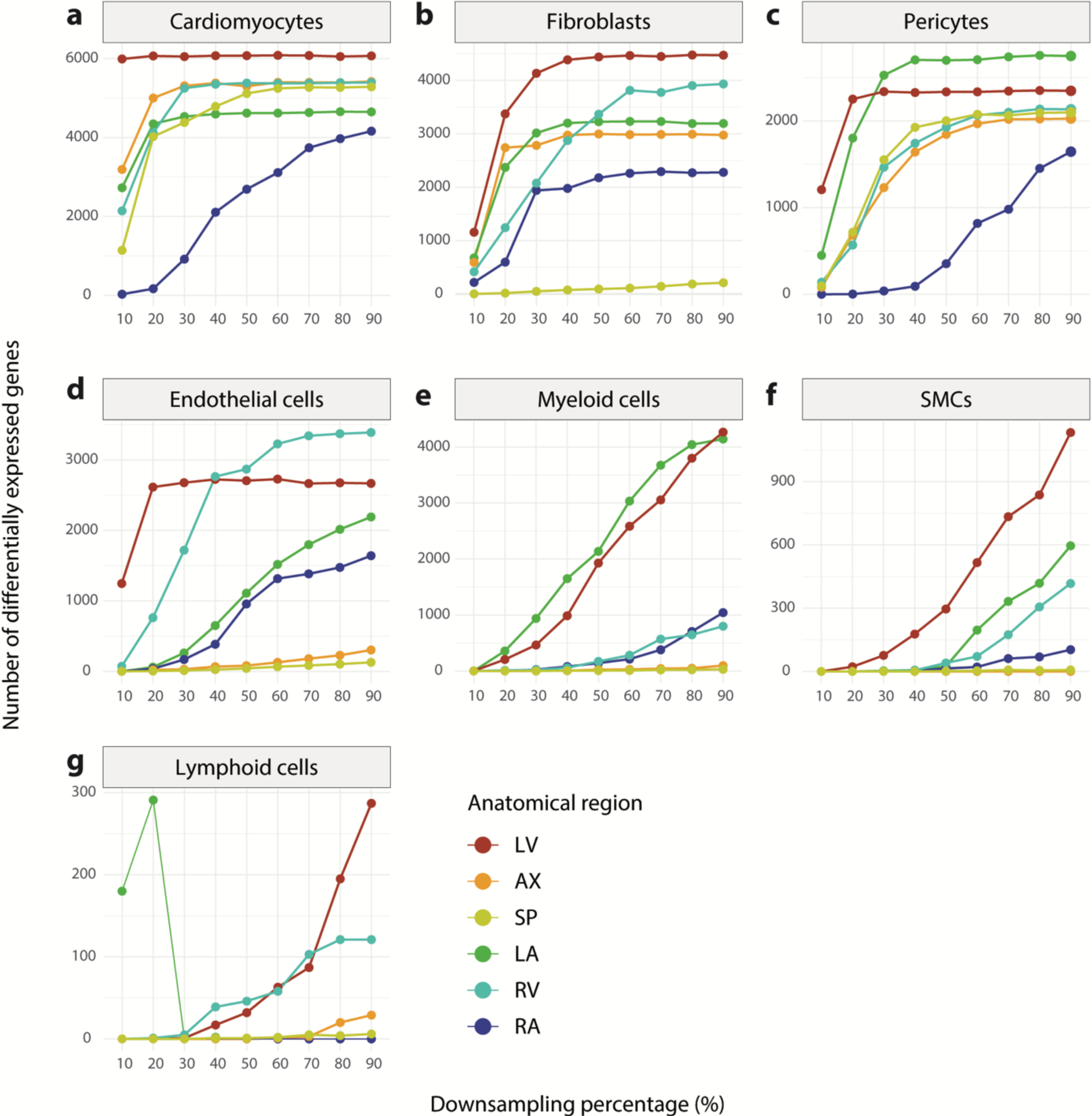
Saturation analysis demonstrates sufficient statistical power to detect sex-biased genes in most regional cell types. Number of significantly (adj. *p* < 0.05) sex-biased genes detected using proportional down samples of original dataset for each regional cell type. LV, left ventricle; AX, apex; SP, septum; LA, left atrium; RV, right ventricle; RA, right atrium.

**Supplementary Figure 7.**
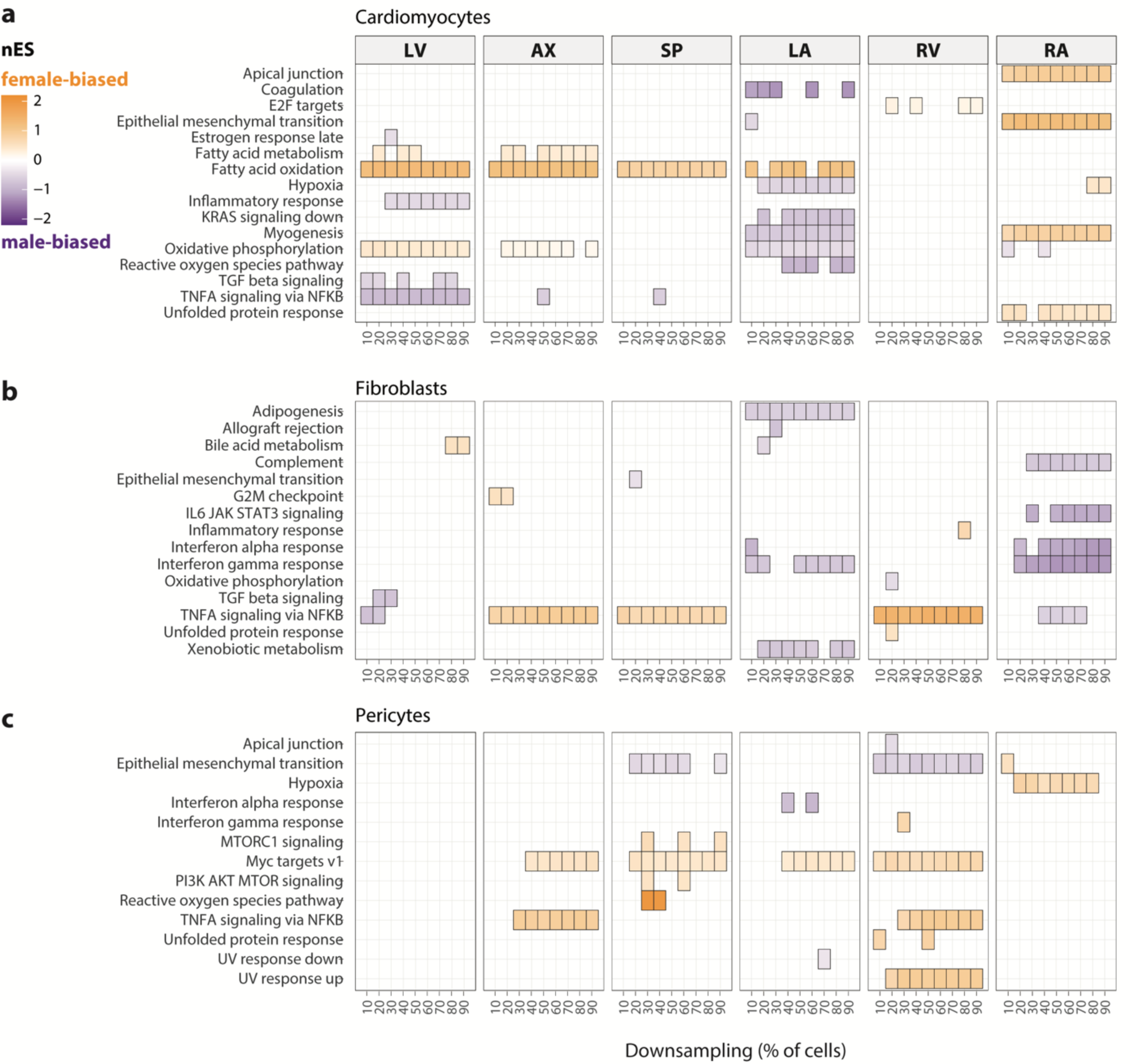
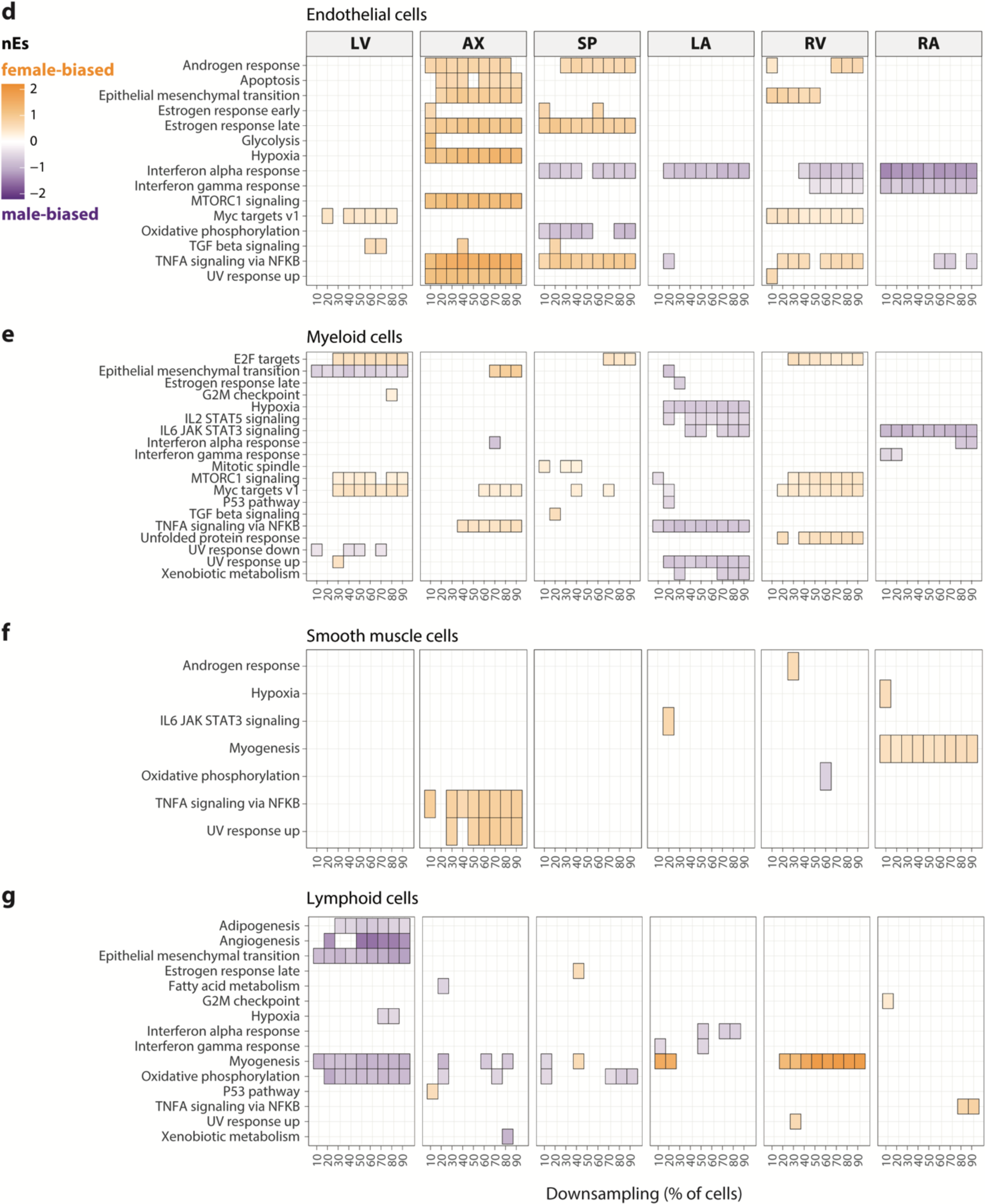
Saturation analysis shows sufficient statistical power to detect sex-biased pathways within the majority of regional cell types. Significantly sex-biased pathways detected using proportional down samples of original dataset for each regional cell type (adj. *p* < 0.05). LV, left ventricle; AX, apex; SP, septum; LA, left atrium; RV, right ventricle; RA, right atrium.

**Supplementary Figure 8.**
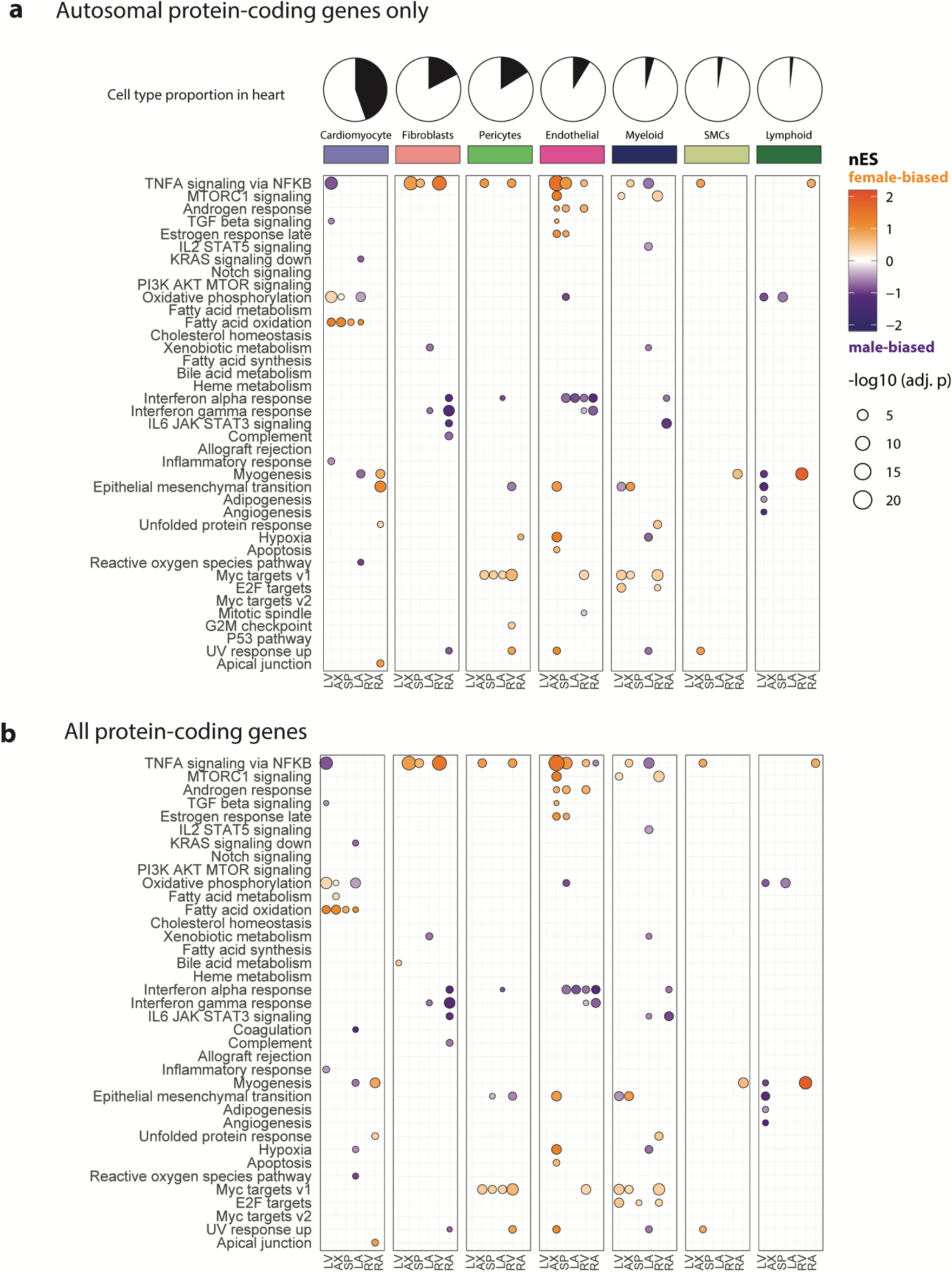
Significantly sex-biased hallmark pathways in cardiac cell types and anatomical regions. Significantly female-biased (orange) and male-biased (purple) Hallmark pathways displayed as a normalized enrichment score (nES) calculated by gene set enrichment analysis with (**a**) autosomal and (**b**) all (autosomal and sex-linked) protein-coding genes. LV, left ventricle; AX, apex; SP, septum; LA, left atrium; RA, right atrium; RV, right ventricle; SMCs, smooth muscle cells.

**Supplementary Figure 9.**
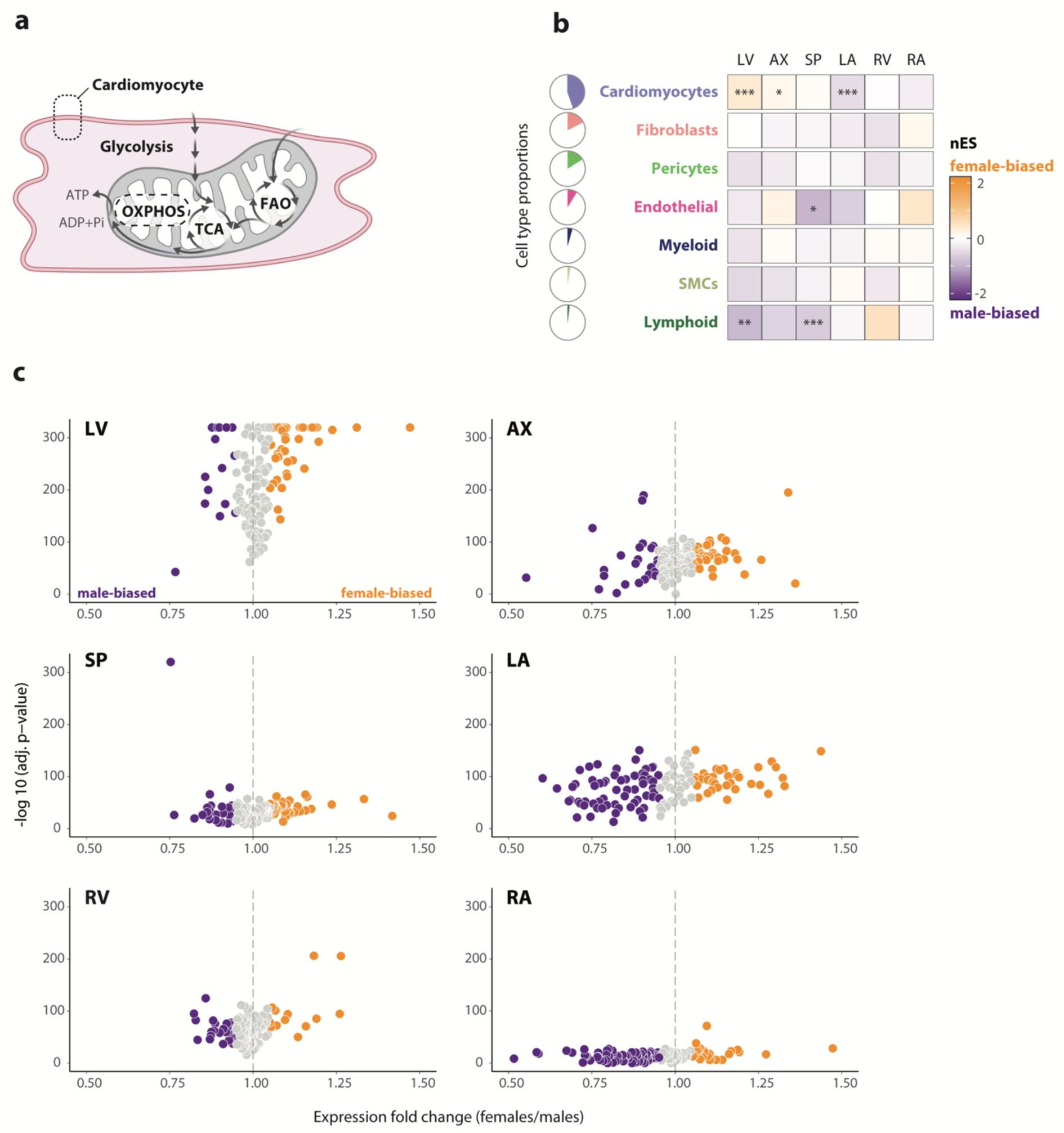
Sex differences in OXPHOS transcript levels. **(a)** Schematic of cardiomyocyte energy metabolism, highlighting the OXPHOS pathway. **(b)** Sex bias of OXPHOS pathway using gene set enrichment analysis across all cardiac cell types and regions. Significantly female-biased (orange) and male-biased (purple) pathways are labeled as follows: adj. *p* < 0.05 (*), adj. *p* < 0.01 (**), adj. *p* < 0.001 (***). **(c)** Volcano plot of significantly male-biased (purple; FC < 0.95, adj. *p* < 0.05) and female-biased (orange, FC > 1.05, adj. *p* < 0.05) OXPHOS genes in cardiomyocytes in each heart anatomical region, with non-significant genes in gray. OXPHOS, oxidative phosphorylation; TCA, tricarboxylic acid cycle; FAO, fatty acid oxidation, LV, left ventricle; AX, apex; SP, septum; LA, left atrium; RV, right ventricle; RA, right atrium; SMCs, smooth muscle cells.

**Supplementary Figure 10.**
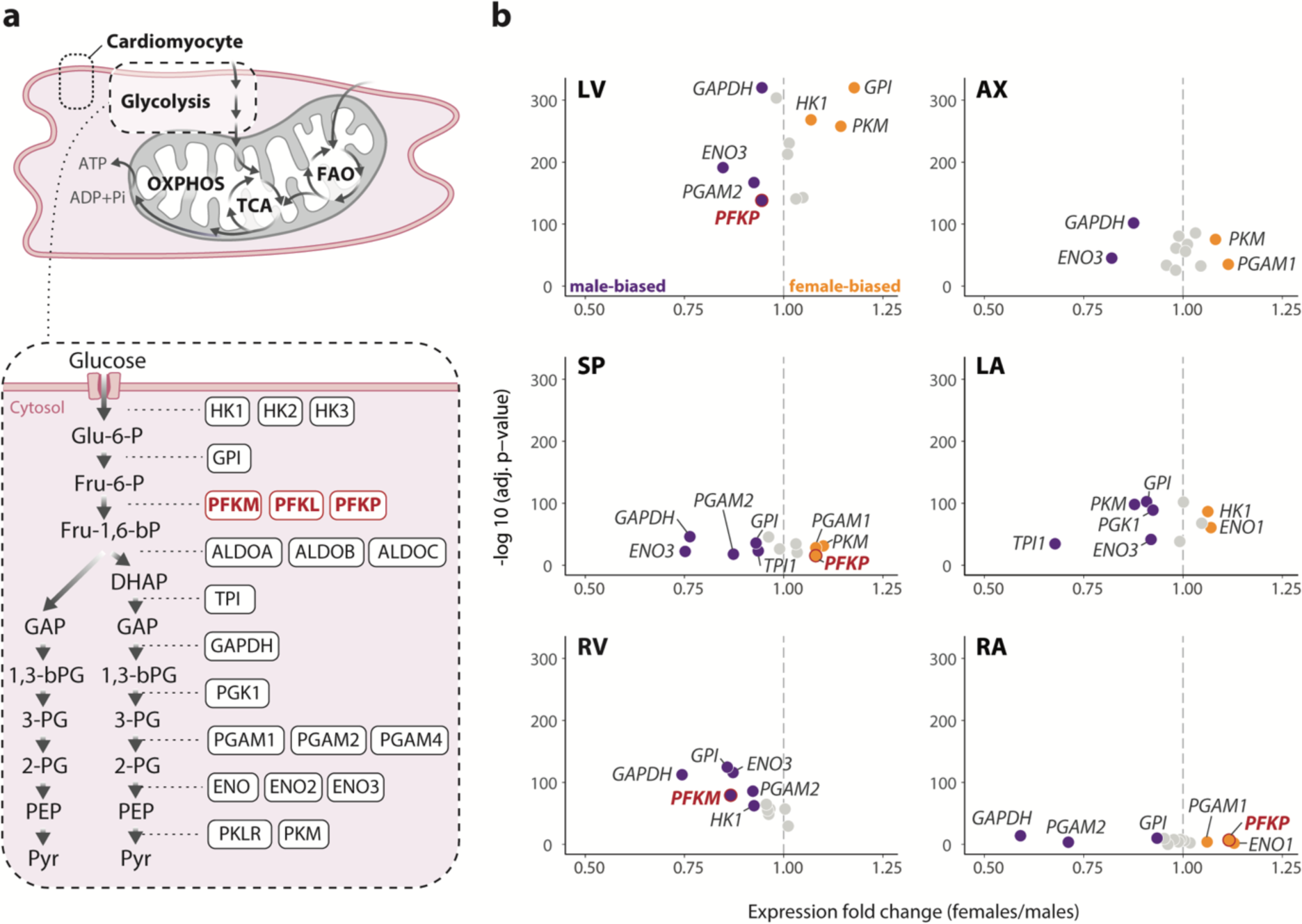
Sex differences in glycolytic transcript levels. **(a)** Schematic of cardiomyocyte energy metabolism, highlighting the glycolytic pathway. Enzymes catalyzing the rate-limiting step of glycolysis in red**. (b)** Volcano plot of significantly male-biased (purple; FC < 0.95, adj. *p* < 0.05) and female-biased (orange, FC > 1.05, adj. *p* < 0.05) glycolytic genes in cardiomyocytes in each heart anatomical region, with non-significant genes in gray. OXPHOS, oxidative phosphorylation; TCA, tricarboxylic acid cycle; FAO, fatty acid oxidation; Glu-6-P, glucose-6-phosphate; Fru-6-P, fructose-6-phosphate; Fru-1,6-bP, fructose-1-3-bisphosphate; DHAP, dihydroxyacetone phosphate; GAP, glyceraldehyde phosphate; 1,3-bPG, 1,3-bisphosphoglycerate; 3-PG, 3-phosphoglycerate; 2-PG, 2-phosphoglycerate; PEP, phosphoenolpyruvate; Pyr, pyruvate; LV, left ventricle; AX, apex; SP, septum; LA, left atrium; RV, right ventricle; RA, right atrium.

**Supplementary Figure 11.**
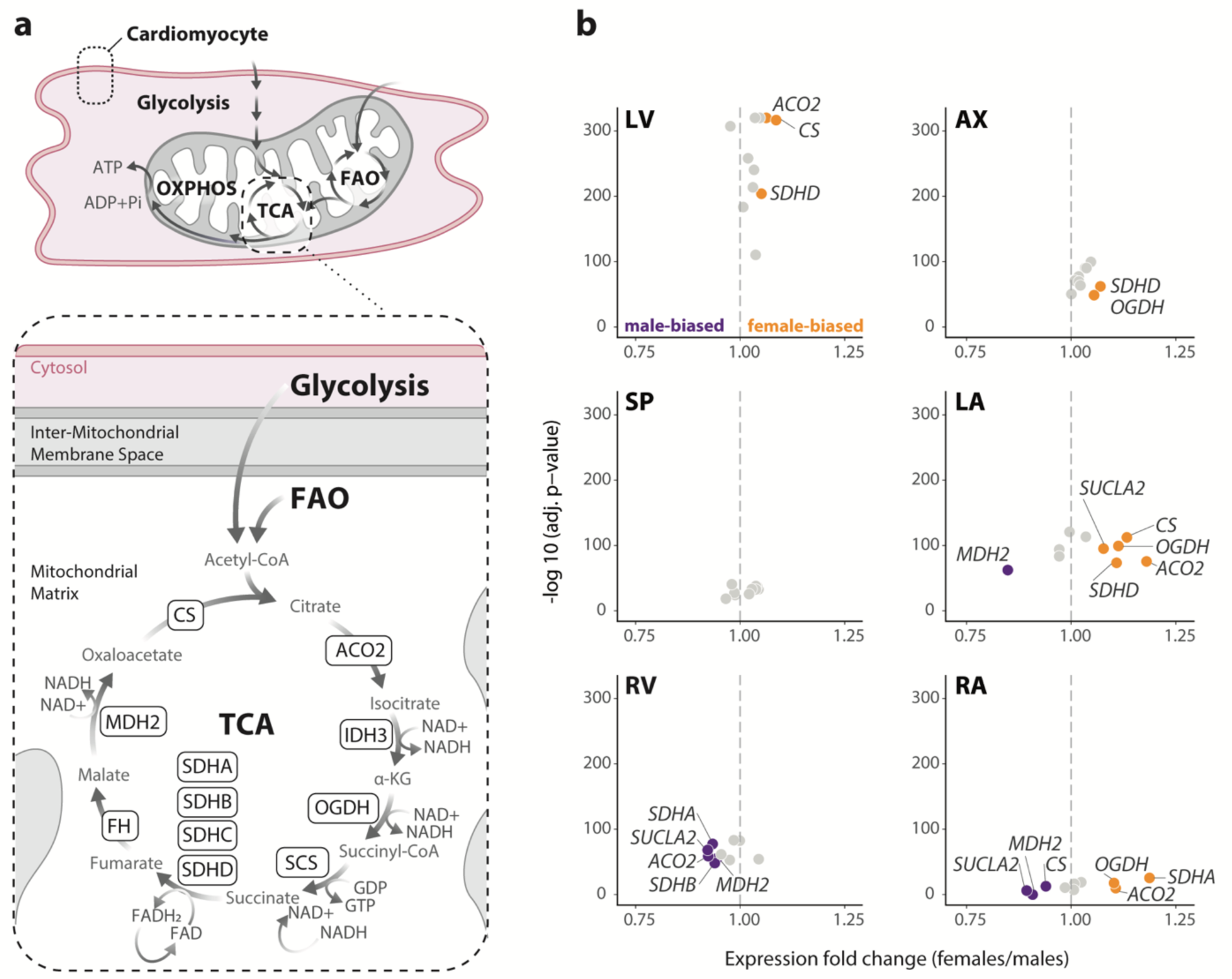
Sex differences in TCA cycle transcript levels. **(a)** Schematic of cardiomyocyte energy metabolism, highlighting the TCA cycle. **(b)** Volcano plot of significantly male-biased (purple; FC < 0.95, adj. *p* < 0.05) and female-biased (orange, FC > 1.05, adj. *p* < 0.05) TCA cycle genes in cardiomyocytes in each heart anatomical region, with non-significant genes in gray. OXPHOS, oxidative phosphorylation; TCA, tricarboxylic acid cycle; FAO, fatty acid oxidation, LV, left ventricle; AX, apex; SP, septum; LA, left atrium; RV, right ventricle; RA, right atrium.

**Supplementary Figure 12.**
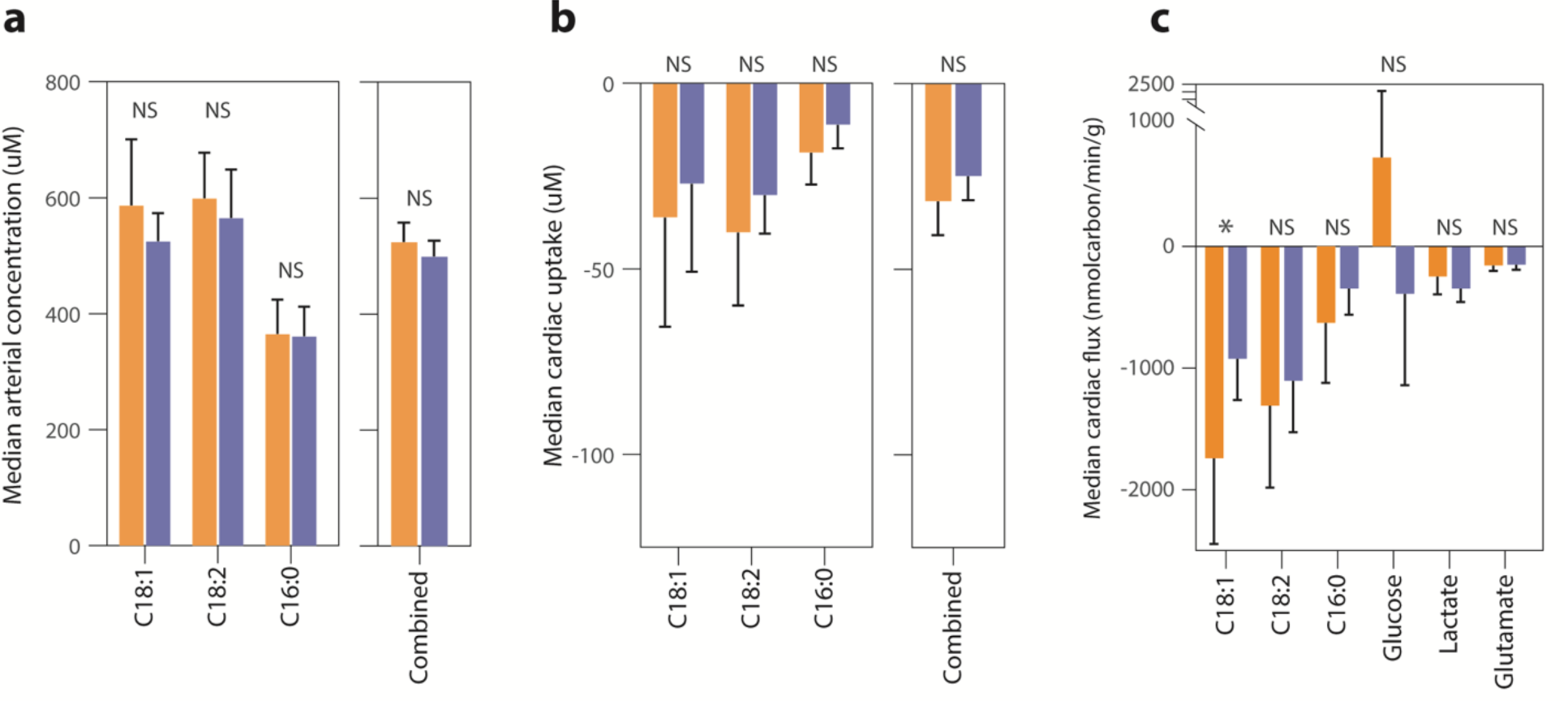
Median arterial concentration, cardiac uptake, and cardiac flux of metabolites in males and females. **(a)** Arterial concentration (median, with 95% confidence interval) of individual (linoleic acid C18:1, oleic acid C18:2, palmitic acid C16:0) or combined free fatty acids (FFAs) shown in female (orange, n = 34) and male individuals (purple, n = 53) from Murashige *et al.,* 2020. **(b)** Cardiac uptake (median, with 95% confidence interval) of individual and combined FFAs from Murashige *et al.* **(c)** Myocardial flux (median, with 95% confidence interval) of C18:1, C18:2 and C16:0 calculated with data from Murashige *et al*. and Ngo *et al.,* 2022. Across all comparisons, the only significant sex difference is in myocardial flux of C18:1 (Welch’s t-test *p* = 0.037). NS, not significant.

**Supplementary Figure 13.**
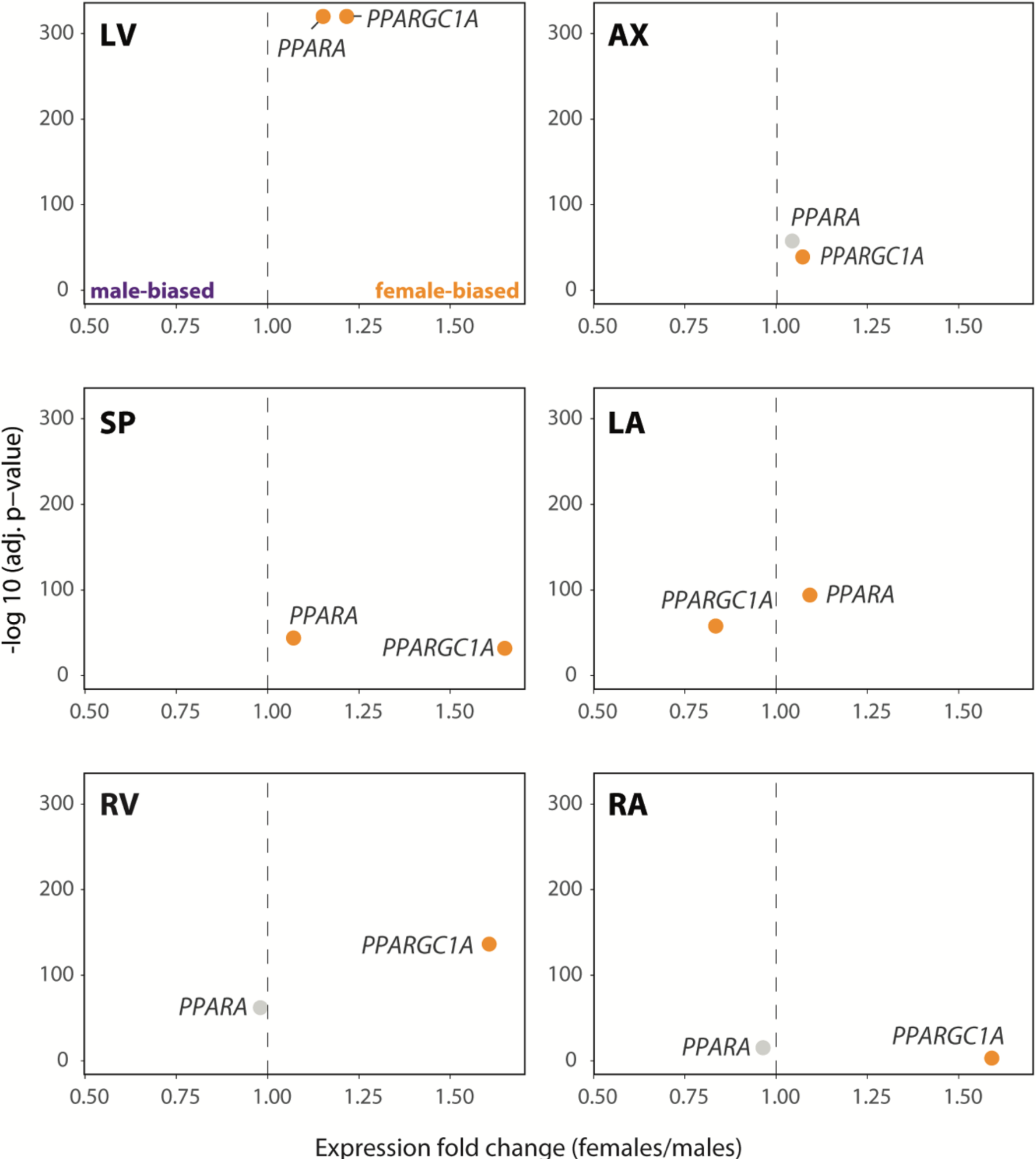
Sex differences in *PPARGC1A* and *PPARA* expression. (**a**) Volcano plot of *PPARGC1A* and *PPARA* expression in cardiomyocytes in each heart anatomical region. Significantly male-biased genes are labeled in purple (FC < 0.95, adj. *p* < 0.05); significantly female-biased genes are labeled in orange (FC > 1.05, adj. *p* < 0.05); non-significant genes in gray. LV, left ventricle; AX, apex; SP, septum; LA, left atrium; RA, right atrium; RV, right ventricle.

